# Tubular lysosome induction couples animal starvation to healthy aging

**DOI:** 10.1101/2021.10.28.466256

**Authors:** Tatiana V. Villalobos, Bhaswati Ghosh, Sanaa Alam, Tyler J. Butsch, Brennan M. Mercola, Cara D. Ramos, Suman Das, Eric D. Eymard, K. Adam Bohnert, Alyssa E. Johnson

## Abstract

Dietary restriction promotes longevity via autophagy activation. However, changes to lysosomes underlying this effect remain unclear. Using the nematode *Caenorhabditis elegans*, we show that induction of autophagic tubular lysosomes, which occurs upon dietary restriction or mTOR inhibition, is a critical event linking reduced food intake to lifespan extension. We find that starvation induces tubular lysosomes not only in affected individuals but also in well-fed descendants, and the presence of gut tubular lysosomes in well-fed progeny is predictive of enhanced lifespan. Furthermore, we demonstrate that expression of *Drosophila* SVIP, a tubular-lysosome activator in flies, artificially induces tubular lysosomes in well-fed worms and improves *C. elegans* health in old age. These findings identify tubular lysosomes as a new class of lysosomes that couples starvation to healthy aging.

**One-Sentence Summary:** Tubular lysosome induction promotes healthy aging.

## Results and Discussion

Dietary restriction (DR) enhances intestinal fitness and extends healthy lifespan of multiple species, in part, by stimulating autophagy (*1–6*). Lysosomes are the terminal site for autophagic degradation and are also major signaling hubs that can sense nutrient scarcity to promote their own biogenesis (*7–10*). Thus, lysosomes possess a remarkable versatility to sense and respond to metabolic shifts in the cell. But whether alternative mechanisms, in addition to increased biogenesis, can modulate lysosome activity in response to nutrient deprivation has not been fully explored. Recently, dynamic autophagic tubular lysosomes (TLs) have been described in *Drosophila* and *C. elegans* (*11–14*). Notably, TLs are cell-type specific and, in some tissues, respond to different metabolic or developmental cues (*12, 13*). Of note, TLs in the gut are robustly stimulated by starvation cues and are naturally induced during early aging to coordinate bulk age-dependent turnover of peroxisomes and potentially other autophagic cargo (*14*). In *Drosophila* larval body-wall muscles, TLs are constitutively present, but increasing their density in muscles alone is sufficient to extend animal lifespan and their disruption leads to multi-system degeneration (*15*). Collectively, these studies allude to TLs as pro-health factors and indicate that, in some tissues, TLs are stimulated by nutrient deprivation. Taken together, we were prompted to determine whether TLs contribute to the known beneficial effects of DR.

In *Drosophila* and *C. elegans*, TLs can be visualized in live tissues by over-expressing a fluorescently-tagged lysosomal membrane protein, Spinster (*11, 14*). To study TLs in their most natural setting, we used CRISPR to insert an mCherry tag at the endogenous C-termini of three *C. elegans* Spinster homologs: *spin-1, spin-2* and *spin-3*. We then examined the native expression pattern of each paralog; *spin-1* is predominantly expressed throughout the intestine and uterus, *spin-2* in the pharynx and *spin-3* predominantly in the posterior region of the intestine (Fig. S1A-C). We focused on *spin-1::mcherry* for our studies since *spin-1* showed the strongest expression pattern in the intestine and intestinal fitness is strongly linked to the beneficial effects of DR (*3*). Upon starvation, we found that intestinal lysosomes labeled by endogenous SPIN-1::mCherry transformed from static vesicles into dynamic tubular networks (Fig. 1A-C and Fig. S2), similar to our previous results using a gut-specific *spin-1::mCherry* transgene (*14*). Strikingly, endogenous SPIN-1::mCherry intensities also increased significantly upon starvation (Fig. 1D-E). Lysosomes labeled by endogenous SPIN-2::mCherry and SPIN-3::mCherry also transformed from vesicles to tubules and increased in intensity upon starvation (Fig. S3A-F). Thus, *spin* expression might be under the control of starvation cues, potentially to promote TL induction. Using *eat-2* mutant animals, which provide a genetic model for DR (*4*), we found that TLs and SPIN-1::mCherry intensity likewise increased in calorically-restricted adults (Fig. 1F-J). Notably, we found that TLs were also induced upon starvation in the *Drosophila* salivary gland (Fig. S4A-C), signifying that starvation-dependent TL induction is a conserved phenomenon across multiple animal species and tissues.

**Fig. 1:**
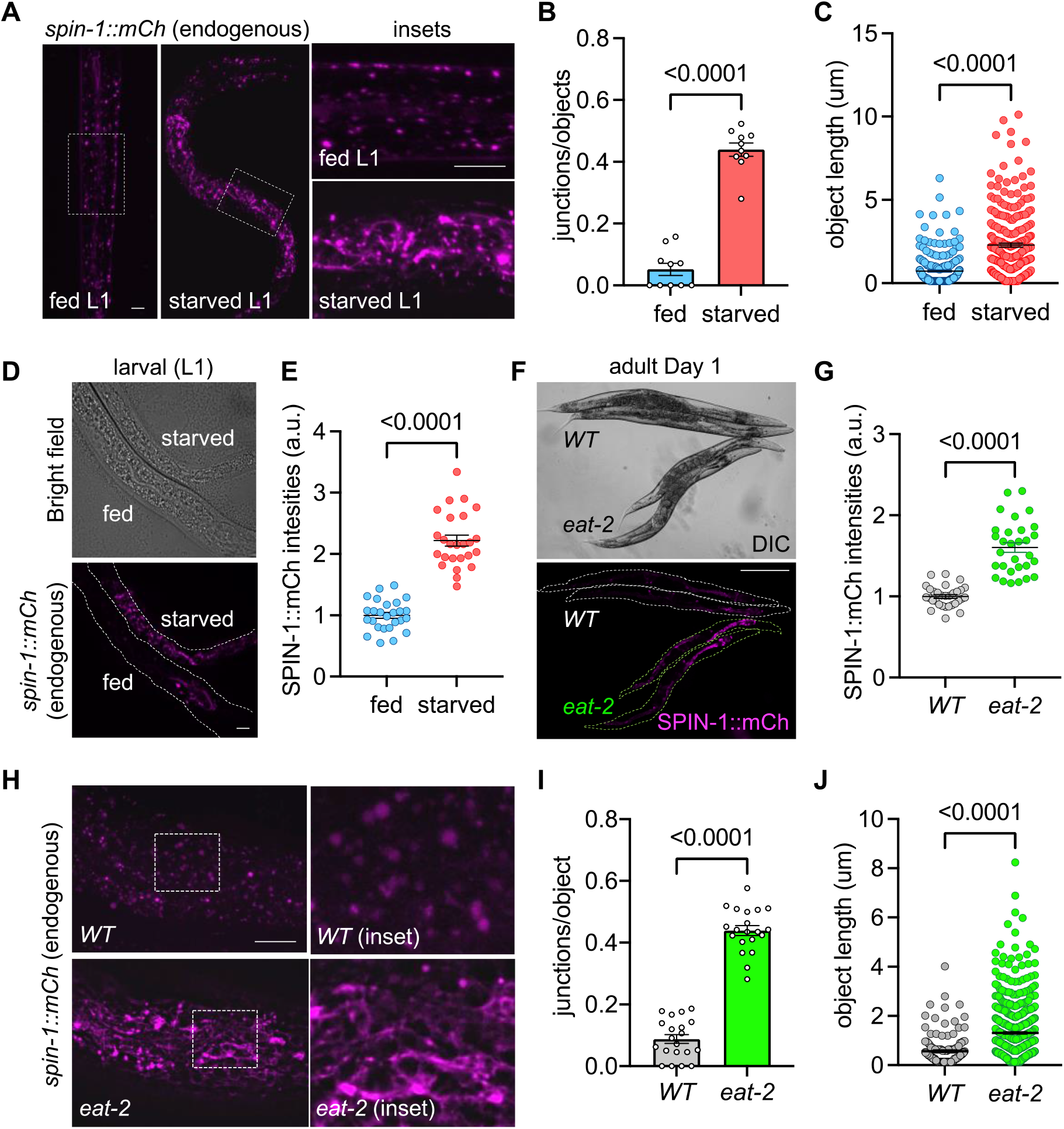
Starvation induces dynamic TLs in the gut. (**A**) Representative images of endogenously-tagged *spin-1::mCh* in fed and starved L1 worms. Bars, 5μm. (**B-C**) Quantification of lysosome junctions/object (B) and length (C) in fed and starved L1 worms. (**D**) Representative images of *spin-1* expression in fed and starved L1 worms. Bars, 5μm. (**E**) Quantification of SPIN-1::mCh intensities in fed and starved L1 worms. (**F**) Representative images of *spin-1* expression in *WT* and *eat-2* day 1 adults. Bar, 100μm. (**G**) Quantification of SPIN-1::mCh intensities in *WT* and *eat-2* day 1 adults. (**H**) Representative images of endogenously-tagged *spin-1::mCh* in *WT* and *eat-2* day 1 adults. Bars, 5μm. (**I-J**) Quantification of lysosome junctions/object (I) and object length (J) in *WT* and *eat-2* day 1 adults. Statistical significance was determined using student’s t-test (p-values indicated on graphs).

Nutrient deprivation is a major autophagic stimulant (*16–18*). Using a tandem mCherry-GFP-LGG-1 autophagy flux reporter, we found that starvation triggered robust autophagosome turnover at TLs (Fig. 2A-B), consistent with our prior observations of starvation-induced turnover of peroxisomes at TLs (*14*). Because starvation-based autophagy induction relies on inhibition of mTOR signaling (*19–21*), we tested whether TL induction was dependent on mTOR inhibition. Indeed, RNAi-mediated knock-down of *let-363* (mTOR) or *daf-15* (RPTOR) induced TLs in the gut of well-fed animals (Fig. 2C-E). Thus, starvation-induced TLs target autophagosomes and are stimulated by mTOR inhibition. Importantly, this is a notable distinction from the lysosome tubules that have been previously described to function during autophagic lysosome reformation (ALR), whereby proto-lysosome tubules emanate from autolysosomes as a mechanism to replenish the pool of functional lysosomes after autophagy terminates (*22*). The proto-lysosome tubules generated via ALR are devoid of autophagic cargo, are not acidic and require mTOR re-activation for their induction (*22*). In contrast, the intestinal TLs we observe are triggered during autophagy to internalize autophagosomes (Fig. 2A-B), are acidic (*14*) and require mTOR inhibition (Fig. 2C-E), rather than activation. Thus, starvation-induced TLs are distinct from ALR and are associated with active autophagy.

**Fig. 2:**
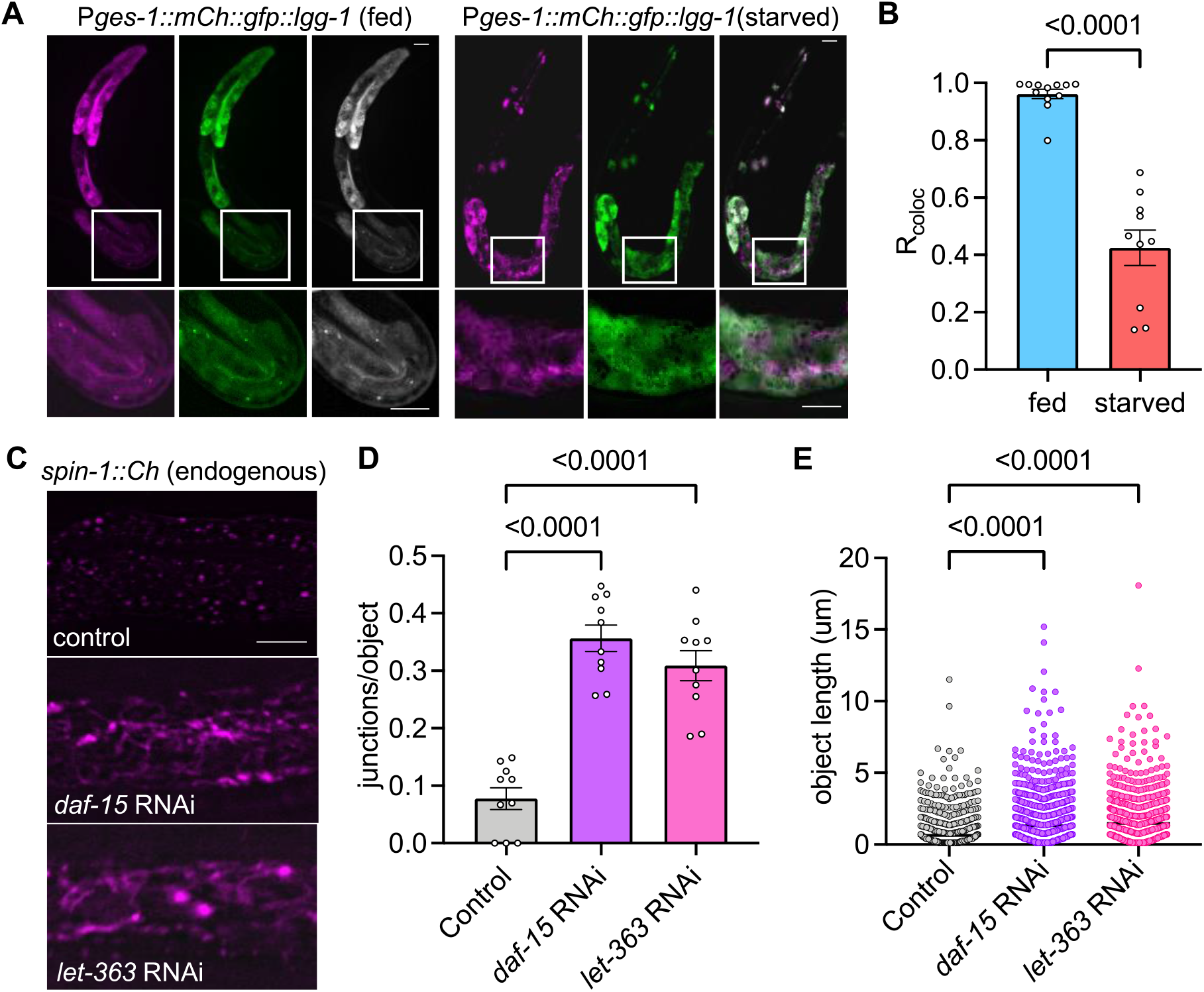
Gut TLs are autophagic and stimulated by TOR inhibition. (**A**) Representative images of *mCh::GFP::lgg-1* expressed in the gut of fed and starved L1 worms. (**B**) Quantification of colocalization between mCh and GFP channels in fed and starved L1 worms. (**C**) Representative images of endogenously tagged *spin-1::mCh* worms fed control, *daf-15* or *let-363* RNAi. (**D-E**) Quantification of lysosomal junctions/object (D) and length (E) in worms fed control, *daf-15* or *let-363* RNAi. Statistical significance was determined using one-way ANOVA with Dunnett’s multiple comparisons (p-values indicated on graphs). Bars, 5μm.

A striking phenomenon of dietary restriction is that the beneficial physiological effects can persist for multiple generations; descendants of starved parents exhibit similar lifespan extension, despite never encountering starvation themselves (*23, 24*). This prompted us to explore whether TLs also persist in subsequent generations of starved parents. Synchronized embryos were seeded onto plates with no food, which induces an L1 larval arrest. After 5 days of prolonged starvation, arrested L1 larvae were transferred to food, which allows their development to resume to adulthood (Fig. 3A). The offspring of these starved parents were then imaged. Remarkably, F1 progeny displayed similar levels of TLs compared to their parents despite being well-fed (Fig. 3B-C). This effect persisted for up to four generations, but the penetrance diminished significantly by the F3 generation (Fig. 3B-C). One mechanism by which starvation elicits transgenerational effects in *C. elegans* is by inducing expression of silencing RNAs (siRNAs), which persist throughout life and transmit epigenetic information to the next generation (*23*). Thus, we tested whether transgenerational induction of TLs is also dependent on siRNAs by examining TLs in the gut of *rde-4* and *hrde-1* mutant worms, which are defective in siRNA production and transmission, respectively (*25, 26*). In both mutants, we observed that TLs were no longer robustly transmitted even in the immediate offspring of starved animals (Fig. 3D-E). Thus, TLs are induced across multiple generations, and this effect is dependent on siRNA transmission.

**Fig. 3:**
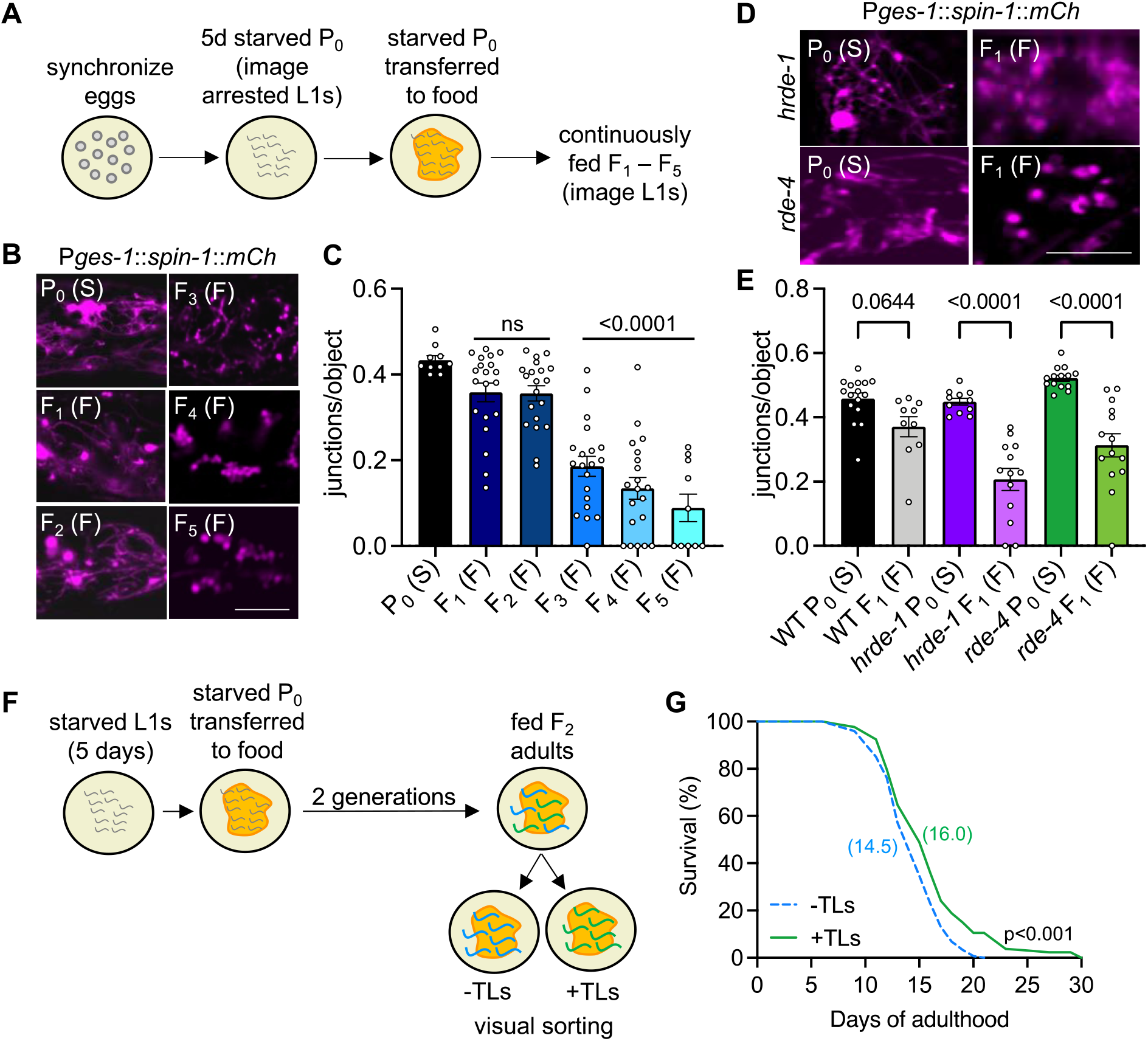
gut TLs are induced transgenerationally via an RNAi mechanism. (**A**) Synchronized eggs were seeded onto NG agar with no food source. After 5 days, starved L1-arrested worms were transferred to food and worm populations were continuously fed and imaged for all subsequent generations. (**B**) Representative images of *spin-1::mCh* expressed in the gut of the indicated generations. (S=Starved, F=Fed). (**C**) Quantification of lysosomal junctions/object from (B). (**D**) Representative images of *spin-1::mCh* expressed in the gut of *hrde-1* and *rde-4* mutants, which are defective in siRNA production and transmission, respectively. (S=Starved, F=Fed). (**E**) Quantification of lysosomal junctions/object from (D). (**F**) Synchronized eggs were seeded onto NG agar with no food source. After 5 days, starved L1-arrested worms were transferred to food and worm populations were continuously fed for 2 generations before sorting. (**G**) Lifespan of progeny from starved grand-parents with (+TLs) or without (-TLs). Mean lifespans are indicated on the graph. For TL analyses, statistical significance was determined using one-way ANOVA with Dunnett’s multiple comparisons. For lifespans, a log-rank test was used to determine statistical significance. p-values are indicated on the graphs. Bars, 5μm.

Because TLs were induced transgenerationally, but with decreasing penetrance in each new generation, (Fig. 3B-C), we wondered whether transgenerational induction of TLs could provide a predictive marker to distinguish the longer-lived individuals in these subsequent populations. To test this, we sorted F2 descendants derived from starved grandparents based on the visual presence or absence of robust TLs in the gut (Fig. 3F). We then compared the lifespan of these sibling populations. Remarkably, we found that F2 worms that retained TLs, lived significantly longer (p<0.001) than their sibling cohorts without TLs (Fig. 3G and S5A-B). Taken together, our data demonstrate that starvation-induced TLs persist for multiple generations and may provide a predictive cellular marker for longer-lived individuals.

Although the lifespan extension observed in animals with TLs is certainly intriguing, this observation is merely correlative and does not directly implicate TLs in promoting animal health and/or longevity since starvation induces multiple metabolic pathways that might contribute to this physiological phenotype. To test whether TLs directly contribute to the pro-health effects of DR, we asked two questions: (1) are TLs required for DR-dependent lifespan extension and (2) are TLs sufficient to exert pro-health effects on their own? To address the first question, we examined whether genetic mutation of the critical TL gene, *spin-1*, could reduce the lifespan extension of *eat-2* mutants. Mutation of *spin-1* alone only slightly reduced the lifespan of *eat-2* mutants (Fig. S6A). However, we considered that multiple *spin* paralogs might have redundant functions and thus tested various combinations of mutant *spin* alleles for their effect on *eat-2* lifespan. We found that *eat-2* lifespan extension was more significantly reduced by double mutation of *spin-1* and *spin-2*, and *eat-2* lifespan was further reduced back to wild type in a *spin-1; spin-2; spin-3* triple-mutant background (Figs. 4A and S6B-C). Importantly, analogous effects on lifespan were not seen in *eat-2+* animals; *spin-1; spin-2* double mutants did not display a reduction in lifespan relative to wild type, and *spin-1; spin-2; spin-3* triple mutants displayed only a modest reduction in lifespan under normal conditions (Figs. 4B and S6B-C). Collectively, these data support the notion that TLs are required for full lifespan extension under DR.

**Fig. 4:**
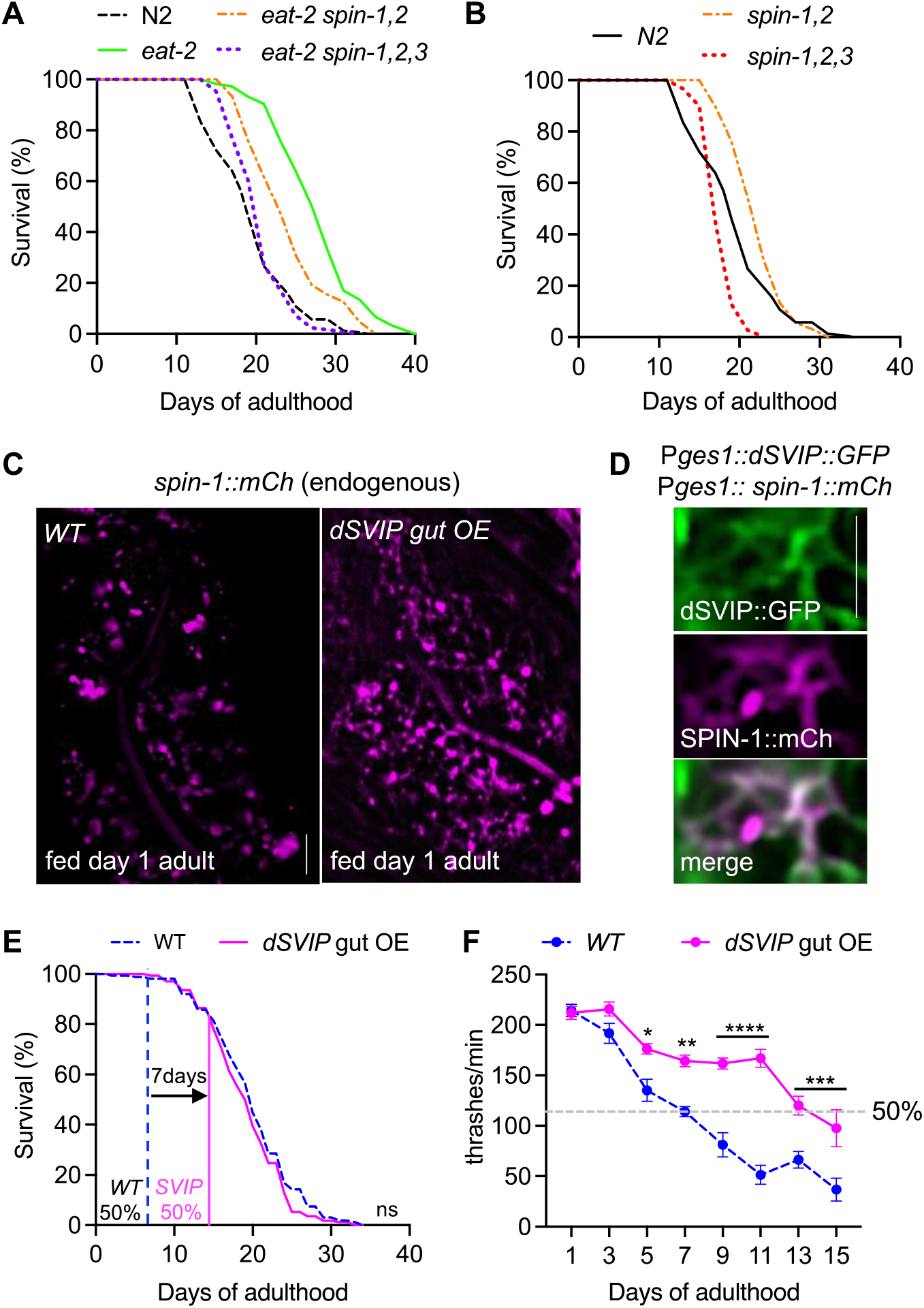
Induction of intestinal TLs promotes healthy aging during DR. (**A-B**) Lifespan of the indicated genotypes. See Fig. S6B-C for mean lifespans and p-values. (**C**) Representative images of endogenously expressed *spin-1::mCh* in fed *WT* worms and worms over-expressing codon-optimized *Drosophila* SVIP (*dSVIP*). (**D**) Co-localization of dSVIP::GFP and SPIN-1::mCh. (**E**) Lifespan of *WT* and *dSVIP* over-expressing worms. Vertical lines indicate the age at which 50% starting thrashing rate was reached for *WT* and *dSVIP* over-expression worms, respectively. (**F**) Thrashing rate of *WT* and *dSVIP* over-expressing worms at the indicated ages. Dashed line represents 50% of starting thrashing rate for both genotypes. Statistical significance was determined using one-way ANOVA with Dunnett’s multiple comparisons (*p<0.05, **p<0.01, ***p<0.001, ****p<0.0001). A log-rank test was used to determine significance for lifespans. Bars, 5μm.

To address whether TLs can exert pro-health effects on their own, we used a genetic strategy to ectopically induce TLs in well-fed animals. Previously, we identified the small VCP-interacting protein (SVIP) as a critical TL modulator; over-expression of *SVIP* in *Drosophila* muscles was sufficient to increase TL density (*15*). The *C. elegans* genome does not possess an annotated *SVIP* ortholog, so we generated a transgenic worm strain to over-express a codon optimized *Drosophila SVIP* gene (*dSVIP*). Indeed, over-expression of *dSVIP* in the *C. elegans* gut was sufficient to induce TL formation under well-fed conditions (Fig. 4C). dSVIP also co-localized with TLs as expected (Fig. 4D). This highlights the general capacity of SVIP to induce TLs artificially in multiple species. We then explored the physiological effects of *dSVIP* over-expression in the *C. elegans* gut. Although we observed no significant effect on lifespan (Fig. 4E), we noticed that *dSVIP* over-expressing worms appeared to retain their mobility later into their life. To investigate this further, we assayed the thrashing rate throughout the lifespan of control and *dSVIP* over-expression worms. Remarkably, *dSVIP* over-expression worms remained more active significantly later into their life (Fig. 4F, Movie S1-S2). In fact, the age at which *dSVIP* over-expression worms reached 50% of their starting thrashing rate was double that of wild type worms (Fig. 4F). Prolonged mobility is a strong indicator of increased healthspan (*27*); thus, while *SVIP*-dependent TL induction in the gut did not extend the average age of mortality, TLs might promote healthier aging.

Taken together, our results demonstrate that, in some tissues, lysosomes can mobilize into highly digestive tubular networks in response to nutrient deprivation to promote organismal health. Significantly, the pro-health effects of TL induction may extend beyond nutrient deprivation. For instance, we have observed that oxidative stress, which at low levels can extend lifespan (*28–32*), also induces TLs in the gut (Fig. S7A-D). TLs are also induced in the *C. elegans* cuticle specifically at molting stages (*12*), in *Drosophila* abdominal muscles during metamorphosis (*13*) and in the *C. elegans* gut during natural aging (Fig. S8A-D, (*14*)). Potentially, in some tissues, TLs could be restrained and only deployed when large pools of cargo need to be degraded en masse to recalibrate cellular homeostasis during adverse metabolic states or major developmental transitions. The mechanism by which TLs are induced remains unknown, but our observation that SPIN protein intensity coincides with TL induction in multiple biological contexts, including starvation, oxidative stress and natural aging (Figs. 1E-F, S7C-D, S8C-D and (*14*)), suggests that expression levels of *spin* genes may be one key determinant of TL induction. However, other molecular factors likely coordinate with *spin* to induce TLs as *spin-1* over-expression alone is not sufficient to induce TLs (Fig. 3B). In sum, we have provided strong evidence that TL induction promotes healthy aging and could be a viable target for age-related disease interventions.

## Acknowledgments

The authors thank all members of the Bohnert and Johnson labs for helpful discussions on this project.

## Author contributions

Conceptualization: KAB, AEJ

Methodology: TVV, EDE, KAB, AEJ

Investigation: TVV, BG, SA, TJB, BMM, CDR, SD, KAB, AEJ

Visualization: KAB, AEJ

Funding acquisition: SA, KAB, AEJ

Project administration: KAB, AEJ

Supervision: KAB, AEJ

Writing – original draft: KAB, AEJ

Writing – review & editing: KAB, AEJ

## Data and materials availability

All data are available in the main text or the supplementary materials.

## Supplementary Materials

Materials and Methods

Figs. S1 to S8

Table S1

Movies S1 to S2

## References and Notes

1. F. Madeo, A. Zimmermann, M. C. Maiuri, G. Kroemer, Essential role for autophagy in life span extension. J. Clin. Invest. 125, 85–93 (2015).

2. M. Hansen, A. Chandra, L. L. Mitic, B. Onken, M. Driscoll, C. Kenyon, A role for autophagy in the extension of lifespan by dietary restriction in C. elegans. PLoS Genet. 4, e24 (2008).

3. K. Akagi, K. A. Wilson, S. D. Katewa, M. Ortega, J. Simons, T. A. Hilsabeck, S. Kapuria, A. Sharma, H. Jasper, P. Kapahi, Dietary restriction improves intestinal cellular fitness to enhance gut barrier function and lifespan in D. melanogaster. PLOS Genet. 14, e1007777 (2018).

4. B. Lakowski, S. Hekimi, The genetics of caloric restriction in Caenorhabditis elegans. Proc. Natl. Acad. Sci. U. S. A. 95, 13091 (1998).

5. F. L, P. L, L. VD, Extending healthy life span--from yeast to humans. Science. 328, 321–326 (2010).

6. K. P, K. M, H. M, Dietary restriction and lifespan: Lessons from invertebrate models. Ageing Res. Rev. 39, 3–14 (2017).

7. C. Settembre, R. Zoncu, D. L. Medina, F. Vetrini, S. Erdin, S. Erdin, T. Huynh, M. Ferron, G. Karsenty, M. C. Vellard, V. Facchinetti, D. M. Sabatini, A. Ballabio, A lysosome-to-nucleus signalling mechanism senses and regulates the lysosome via mTOR and TFEB. EMBO J. 31, 1095–1108 (2012).

8. A. Ballabio, J. S. Bonifacino, Lysosomes as dynamic regulators of cell and organismal homeostasis. Nat. Rev. Mol. Cell Biol. 21 (2020), pp. 101–118.

9. Y. Sancak, L. Bar-Peled, R. Zoncu, A. L. Markhard, S. Nada, D. M. Sabatini, Ragulator-rag complex targets mTORC1 to the lysosomal surface and is necessary for its activation by amino acids. Cell. 141, 290–303 (2010).

10. R. Zoncu, L. Bar-Peled, A. Efeyan, S. Wang, Y. Sancak, D. M. Sabatini, mTORC1 senses lysosomal amino acids through an inside-out mechanism that requires the vacuolar H+-ATPase. Science (80-.). 334, 678–683 (2011).

11. A. E. Johnson, H. Shu, A. G. Hauswirth, A. Tong, G. W. Davis, VCP-dependent muscle degeneration is linked to defects in a dynamic tubular lysosomal network in vivo. Elife. 4 (2015), doi:10.7554/eLife.07366.

12. R. Miao, M. Li, Q. Zhang, C. Yang, X. Wang, An ECM-to-Nucleus Signaling Pathway Activates Lysosomes for C. elegans Larval Development. Dev. Cell. 52, 21–37.e5 (2020).

13. T. Murakawa, A. A. Kiger, Y. Sakamaki, M. Fukuda, N. Fujita, An autophagy-dependent tubular lysosomal network synchronizes degradative activity required for muscle remodeling. J. Cell Sci. 133 (2020), doi:10.1242/jcs.248336.

14. D. A. Dolese, M. P. Junot, B. Ghosh, T. J. Butsch, A. E. Johnson, K. A. Bohnert, Degradative tubular lysosomes link pexophagy to starvation and early aging in C. elegans. https://doi.org/10.1080/15548627.2021.1990647 (2021), doi:10.1080/15548627.2021.1990647.

15. A. E. Johnson, B. O. Orr, R. D. Fetter, A. J. Moughamian, L. A. Primeaux, E. G. Geier, J. S. Yokoyama, B. L. Miller, G. W. Davis, SVIP is a molecular determinant of lysosomal dynamic stability, neurodegeneration and lifespan. Nat. Commun. 12 (2021), doi:10.1038/s41467-020-20796-8.

16. K. Takeshige, M. Baba, S. Tsuboi, T. Noda, Y. Ohsumi, Autophagy in yeast demonstrated with proteinase-deficient mutants and conditions for its induction. J. Cell Biol. 119, 301–312 (1992).

17. N. Mizushima, A. Yamamoto, M. Matsui, T. Yoshimori, Y. Ohsumi, In vivo analysis of autophagy in response to nutrient starvation using transgenic mice expressing a fluorescent autophagosome marker. Mol. Biol. Cell. 15, 1101–11 (2004).

18. J. Kaur, J. Debnath, Autophagy at the crossroads of catabolism and anabolism. Nat. Rev. Mol. Cell Biol. 16, 461–472 (2015).

19. K. Y, F. T, S. T, N. K, O. M, O. Y, Tor-mediated induction of autophagy via an Apg1 protein kinase complex. J. Cell Biol. 150, 1507–1513 (2000).

20. W. S, L. R, H. MN, TOR signaling in growth and metabolism. Cell. 124, 471–484 (2006).

21. S. RC, S. O, N. TP, Role and regulation of starvation-induced autophagy in the Drosophila fat body. Dev. Cell. 7, 167–178 (2004).

22. L. Yu, C. K. McPhee, L. Zheng, G. A. Mardones, Y. Rong, J. Peng, N. Mi, Y. Zhao, Z. Liu, F. Wan, D. W. Hailey, V. Oorschot, J. Klumperman, E. H. Baehrecke, M. J. Lenardo, Termination of autophagy and reformation of lysosomes regulated by mTOR. Nature. 465, 942–6 (2010).

23. O. Rechavi, L. Houri-Ze’Evi, S. Anava, W. S. S. Goh, S. Y. Kerk, G. J. Hannon, O. Hobert, Starvation-induced transgenerational inheritance of small RNAs in C. elegans. Cell. 158, 277–287 (2014).

24. O. Rechavi, I. Lev, Principles of Transgenerational Small RNA Inheritance in Caenorhabditis elegans. Curr. Biol. 27, R720–R730 (2017).

25. H. Tabara, E. Yigit, H. Siomi, C. C. Mello, The dsRNA binding protein RDE-4 interacts with RDE-1, DCR-1, and a DExH-Box helicase to direct RNAi in C. elegans. Cell (2002), doi:10.1016/S0092-8674(02)00793-6.

26. B. A. Buckley, K. B. Burkhart, S. G. Gu, G. Spracklin, A. Kershner, H. Fritz, J. Kimble, A. Fire, S. Kennedy, A nuclear Argonaute promotes multigenerational epigenetic inheritance and germline immortality. Nature (2012), doi:10.1038/nature11352.

27. J. H. Hahm, S. Kim, R. Diloreto, C. Shi, S. J. V. Lee, C. T. Murphy, H. G. Nam, C. elegans maximum velocity correlates with healthspan and is maintained in worms with an insulin receptor mutation. Nat. Commun. 6 (2015), doi:10.1038/ncomms9919.

28. J. Lapointe, S. Hekimi, Early mitochondrial dysfunction in long-lived Mclk1+/-mice. J. Biol. Chem. 283, 26217–26227 (2008).

29. J. M. Van Raamsdonk, S. Hekimi, Deletion of the mitochondrial superoxide dismutase sod-2 extends lifespan in Caenorhabditis elegans. PLoS Genet. 5 (2009), doi:10.1371/journal.pgen.1000361.

30. J. M. Copeland, J. Cho, T. Lo, J. H. Hur, S. Bahadorani, T. Arabyan, J. Rabie, J. Soh, D. W. Walker, Extension of Drosophila Life Span by RNAi of the Mitochondrial Respiratory Chain. Curr. Biol. 19, 1591–1598 (2009).

31. S. J. Lee, A. B. Hwang, C. Kenyon, Inhibition of respiration extends C. elegans life span via reactive oxygen species that increase HIF-1 activity. Curr. Biol. 20, 2131–2136 (2010).

32. W. Yang, S. Hekimi, A mitochondrial superoxide signal triggers increased longevity in caenorhabditis elegans. PLoS Biol. 8 (2010), doi:10.1371/journal.pbio.1000556.

