## Supplemental Material for "Tubular lysosome induction couples animal starvation to healthy aging"

**This PDF file includes:**

Materials and Methods
Figs. S1 to S8
Table S1
Captions for Movies S1 to S2

**Other Supplementary Materials for this manuscript include the following:**

Movies S1 to S2

### Materials and Methods

Strain generation

Supplemental Table 1 provides a complete list of strains used in this study. Endogenously-tagged *spin-1::mCherry*, *spin-2::mCherry* and *spin-3::mCherry* were generated by In Vivo Biosystems using CRISPR technology. All other strains used in this study were generated using standard genetic crosses or microinjection (Evans 2006). For genetic crosses, transgenes expressing fluorescent proteins were tracked by stereomicroscopy, and gene deletions and mutations were verified by nested PCRs and/or sequencing. For microinjection, constructs were injected individually or in combination into the gonad of adult N2 hermaphrodites, each at a concentration of 25 ng/µl. Integration of transgenes was achieved using UV irradiation, followed by >5 generations of outcrossing. The following transgenes were generated in-lab:

***Pges-1::mCherry-gfp::lgg-1::unc-54 UTR***

*lgg-1* was PCR-amplified from *C. elegans* genomic DNA with a stop codon and a 5' SacI site using the following primers: Forward, 5'- GGG GAC AAG TTT GTA CAA AAA AGC AGG CTC AAA ATC TAG AGC GAG CTC ATG AAG TGG GCT TAC AAG GAG GAG -3' -3'; Reverse, 5'- GGG GAC CAC TTT GTA CAA GAA AGC TGG GTT TAT TCC TTC TTT TCG ACC TCT CCT CC -3'. This *lgg-1* sequence was then cloned into the pDONR221 Gateway entry vector using BP clonase, and the insert was verified by DNA sequencing. An mCherry-GFP coding sequence was PCR-amplified (from a lab stock plasmid; Dolese et al. paper) without a stop codon but with 5' and 3' SacI sites using the following primers: Forward, 5'- CGA GCT CAT GGT CTC AAA GGG TGA AGA AGA TAA C -3'; Reverse, 5'- CGA GCT CTT TGT ATA GTT CAT CCA TGC CAT GTG TAA TCC -3'. This mCherry-GFP coding sequence was then inserted into pDONR221 *lgg-1* using standard restriction-enzyme cloning at the SacI site, and pDONR221 *mCherry-gfp::lgg-1* was sequence-verified. pDONR221 *mCherry-gfp::lgg-1* was then combined with lab-stock plasmids pDONR P4-P1r P*ges-1* and pDONR P2R-P3 *unc-54* 3' UTR into the pDEST R4-R3 Gateway destination vector using LR clonase.

***Pges-1::svip::unc-54 UTR* and *Pges-1::svip::gfp::unc-54 UTR***

The coding sequence for *Drosophila SVIP* was codon-optimized for *C. elegans* expression using the *C. elegans* Codon Adaptor (S *et al.* 2011). This sequence was then synthesized as a gBlock by Integrated DNA Technologies and PCR-amplified using the following primers: Forward, 5'- GGG GAC AAG TTT GTA CAA AAA AGC AGG CTC AAA AAA AAA TGG GAG CCT GTT TAT CAT GTT GC -3'; Reverse, 5'- GGG GAC CAC TTT GTA CAA GAA AGC TGG GTT TAA GAG GTT TGC CAA CGA AGG TTA G -3'. This *svip* sequence was then cloned into the pDONR221 Gateway entry vector using BP clonase, and the insert was verified by DNA sequencing. To generate a GFP-tagged version, a Gibson-assembly reaction was performed to insert a GFP coding sequence before the *svip* stop codon. The GFP coding sequence was PCR-amplified using the following primers: Forward, 5'- ATG AGT AAA GGA GAA GAA CTT TTC AC -3'; Reverse, 5'- CTA TTT GTA TAG TTC ATC CAT GCC ATG -3'. The plasmid was PCR-amplified as a linear piece of DNA with appropriate overlaps using the following primers: Forward, 5'- CAT GGC ATG GAT GAA CTA TAC AAA TAG ACC CAG CTT TCT TGT ACA AAG TTG -3'; Reverse, 5'- GTG AAA AGT TCT TCT CCT TTA CTC ATA GAG GTT TGC CAA CGA AGG TTA GAT TG -3'. Again, the full insert was verified by DNA sequencing. Ultimately, pDONR221 *svip* and pDONR221 *svip::gfp* were each combined with lab-stock plasmids pDONR P4-P1r P*ges-1* and pDONR P2R-P3 *unc-54* 3' UTR into the pDEST R4-R3 Gateway destination vector using LR clonase.

Animal maintenance

Worms were raised at 20ºC on NGM agar (51.3 mM NaCl, 0.25% peptone, 1.7% agar, 1 mM $CaCl_2$, 1 mM $MgSO_4$, 25 mM $KPO_4$, 12.9 µM cholesterol, pH 6.0). Fed worms were maintained on NGM agar plates that had been seeded with *E. coli* OP50 bacteria.
Synchronous populations of worms were obtained by bleaching young-adult hermaphrodites. Briefly, adult hermaphrodites were vortexed in 1 mL bleaching solution (0.5 M NaOH, 20% bleach) for 5 minutes to isolate eggs, and eggs were then washed three times in M9 buffer (22 mM $KH_2PO_4$, 42 mM $Na_2HPO_4$, 85.5 mM NaCl, 1 mM $MgSO_4$) before plating.

To obtain starved L1 animals, bleached eggs were spotted on NGM agar that lacked OP50 bacteria, and plates were maintained at 20ºC for 18-24 hours before imaging, except where otherwise noted. For aging experiments, synchronous populations of animals were established by bleaching adult hermaphrodites or by synchronous egg-laying. In all aging experiments excluding lifespan analyses (see below), adult worms were picked onto fresh OP50-seeded NGM plates every 1-2 days to maintain adults separate from progeny.

RNAi experiments

RNAi clones were generated for *let-363* and *daf-15* by using Gibson cloning to insert gene fragments into the cut PstI site of the L4440 vector. The *let-363* gene fragment used in Gibson assembly was PCR-amplified from *C. elegans* genomic DNA using the following primers: Forward, 5'- CCA CGT GAC GCG TGG ATC CCC CGG GCT GCA ATG CTC CAA CAA CAC GGA ATT AGT TTT C -3'; Reverse, 5'- CGG TAT CGA TAA GCT TGA TAT CGA ATT CCG AAT GCT GTC GGT GTG GCC AGT GCG AGC TC -3'. The *daf-15* gene fragment used in Gibson assembly was PCR-amplified from *C. elegans* genomic DNA using the following primers: Forward, 5'- CCA CGT GAC GCG TGG ATC CCC CGG GCT GCA ATG GAA GAG GAT AGG AGT ATA ACA CCG -3'; Reverse, 5'- CGG TAT CGA TAA GCT TGA TAT CGA ATT CCG CCA GGA TAT TTT CTG ATC TTC TCC AAA TC -3'. All clones were verified by DNA sequencing. For RNAi experiments, synchronous populations of animals were grown on OP50-seeded NGM plates until late L4 or day 1 of adulthood, at which time they were transferred to RNAi plates (NGM plus 100 ng/µl carbenicillin and 1 mM IPTG) that had been seeded with bacteria expressing the relevant RNAi clone. An empty L4440 vector was used as a negative control.

Paraquat treatment

For paraquat treatment, a 1 M stock solution of paraquat (Acros Organics, AC22732) was made in water and diluted to a final concentration of 0.25 mM (low dose) or 5 mM (high dose) in NGM agar and in the OP50 bacteria seeded onto the plates. Worms were synchronized by bleaching, and late L4 or day 1 adults were transferred to control or paraquat plates. Imaging was performed 1-2 days after plating on control or paraquat plates.

Lifespan analysis

Synchronous populations of animals were transferred as late L4s to NGM plates seeded with OP50 bacteria. Plates were previously spotted with 5 mM FUdR (Acros Organics) to prevent progeny production. >250 animals were analyzed per condition. Animals that exploded, bagged,

or crawled off plates were censored during analysis. Lifespans were analyzed using OASIS 2 software (Han et al. 2016), and statistical significance was assessed using a log-rank test.
For TL sorting lifespan, synchronized eggs were seeded onto NG agar with no food source. After 5 days, starved L1-arrested worms were transferred to food and worm populations were continuously fed for 2 generations. Late L4 F2 progeny from starved grandparents were sorted under a fluorescence stereo microscope and separated into two populations based on visible presences or absence of TLs. Lifespan of the two populations were analyzed using the method described above.

Thrashing assay
Synchronous populations of animals were transferred as late L4s to NGM plates seeded with OP50 bacteria with no FUDR. Worms were transferred to fresh plates every 1-2 days to separate adults from their progeny. To score thrashing rates, individual worms were transferred into a drop of M9 buffer on an NGM plate and the number of thrashes were counted in a 1-min period.

Microscopy
4% agarose (Fisher Bioreagents) pads were dried on a Kimwipe (Kimtech) and then placed on top of a Gold Seal™ glass microscope slide (ThermoFisher Scientific). A small volume of 2 mM levamisole (Acros Organics) was spotted on the agarose pad. Worms were transferred to the levamisole spot, and a glass cover slip (Fisher Scientific) was placed on top to complete the mounting. Live-animal fluorescence microscopy was performed using a Leica DMi8 THUNDER imager, equipped with 10X (NA 0.32), 40X (NA 1.30), and 100X (NA 1.40) objectives and GFP and Texas Red filter sets.

Image analysis
Images were processed using LAS X software (Leica) and FIJI/ImageJ (NIH). Lysosome networks were analyzed using “Skeleton” analysis plugins in FIJI. Briefly, images were converted to binary 8-bit images and then to skeleton images using the “Skeletonize” plugin. Skeleton images were then quantified using the “Analyze Skeleton” plugin. Number of objects, number of junctions and object lengths were scored. An “object” is defined by the Analyze Skeleton plugin as a branch connecting two endpoints, an endpoint and junction or two junctions. Junctions/object was used as a parameter to quantify network integrity. Correlation coefficients for red fluorescence relative to green fluorescence were quantified by selecting a region of interest in FIJI/ImageJ and using the Coloc 2 plug-in with a point-spread function of 3.0.

Statistical analyses
Data were statistically analyzed using GraphPad Prism. For two sample comparisons, an unpaired t-test was used to determine significance ($\alpha$=0.05). For three or more samples, a one-way ANOVA with Dunnett’s multiple comparisons was used to determine significance ($\alpha$=0.05). Statistical significance of lifespan data was determined using a Log-rank test.

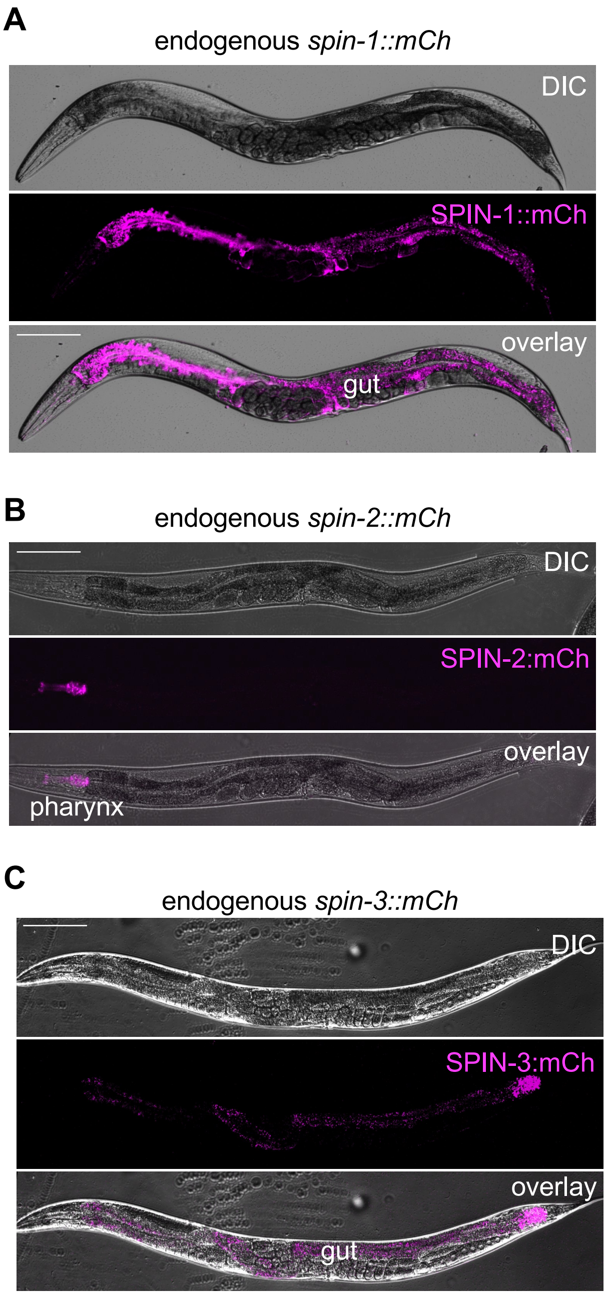


**Fig. S1, related to Fig. 1: *spin* isoform expression patterns**
(**A-C**) Endogenous expression pattern of *spin-1::mCh* (A), *spin-2::mCh* (B) and *spin-3::mCh* (C) in adult worms. Bars, 100µm.

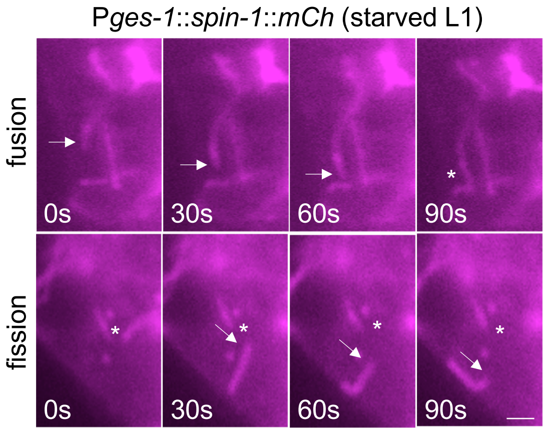


**Fig. S2, related to Fig. 1: TL dynamics in starved animals.**
Representative time-lapse images of SPIN-1::mCh labeled TLs in starved L1 animals demonstrating a fusion (top panel) and fission event (bottom panel). Bar, 2μm.

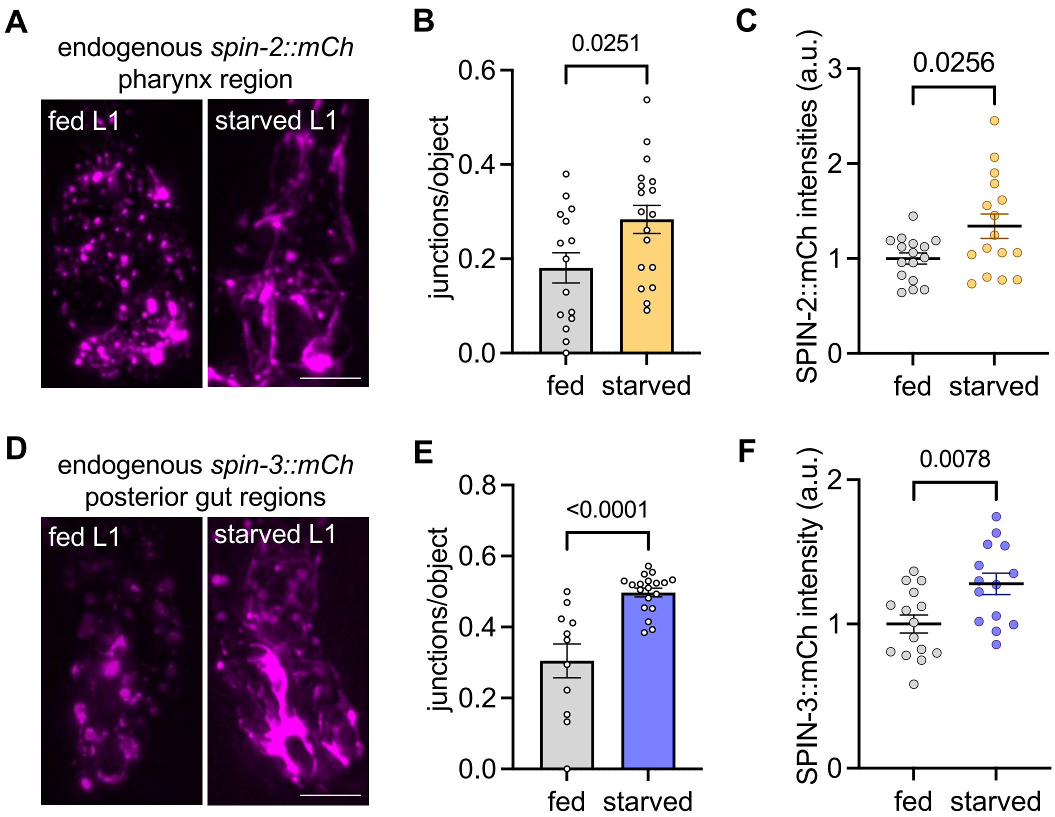


**Fig. S3, related to Fig. 1: SPIN-2 and SPIN-3 localization in fed and starved animals.** (**A**) Representative image of endogenously-tagged *spin-2::mCh* in fed and starved L1 worms. (**B-C**) Quantification of lysosome junctions/object (B) and SPIN-2::mCh intensities (C) in fed and starved L1 worms. (**D**) Representative image of endogenously-tagged *spin-3::mCh* in fed and starved L1 worms. (**E-F**) Quantification of lysosome junctions/object (E) and SPIN-3::mCh intensities (F) in fed and starved L1 worms. Bars, 5µm.

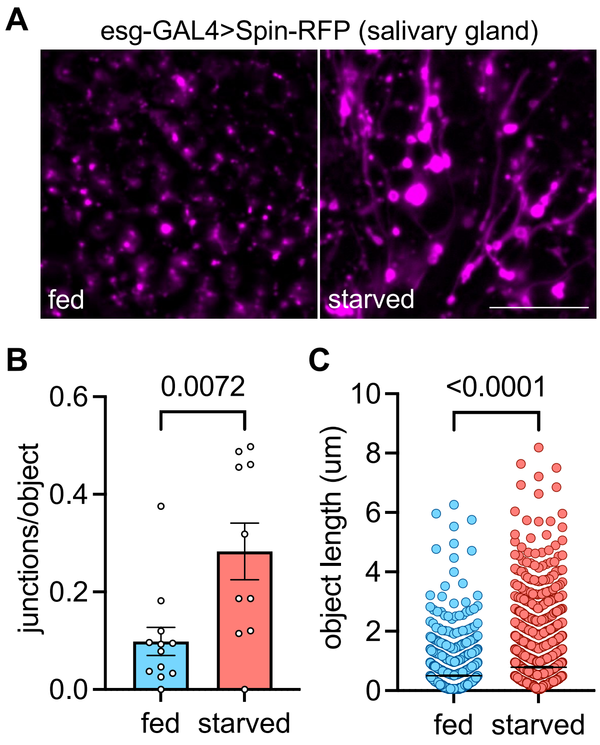


**Fig. S4, related to Fig. 1: Starvation induces TLs in the *Drosophila* salivary gland.**
(**A**) Representative images of Spin-RFP expressed in the salivary gland of in fed and starved *Drosophila* 3rd instar larvae. (**B**) Quantification of lysosomal junctions/object (B) and object length (C) in the salivary gland of in fed and starved *Drosophila* 3rd instar larvae. Bar, 5µm.

**A**

| genotype | n | mean (days) | SEM |
|---|---|---|---|
| -TLs | 147 | 14.46 | 0.22 |
| +TLs | 133 | 15.9 | 0.36 |

**B**

| condition | χ2 | p-value | Corrected p-value |
|---|---|---|---|
| -TLs v.s. +TLs | 13.02 | 0.0003 | 0.0003 |

**Fig. S5, related to Fig. 3: Statistical analyses of TL correlation lifespan.**
(**A**) Descriptive statistics for lifespans in Figure 3G. (**B**) Log-rank test results for lifespan comparisons in Figure 3G.

A

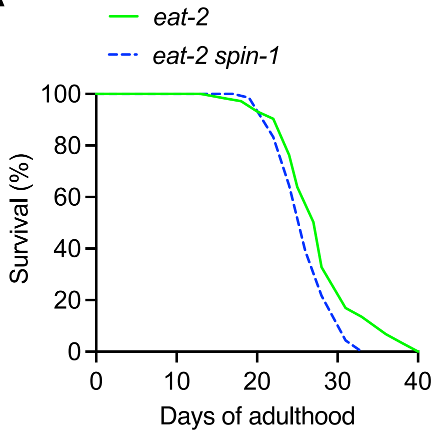


B

| genotype | n | mean (days) | SEM |
|---|---|---|---|
| *Figure S6A* | | | |
| *eat-2* | 177 | 28.06 | 0.38 |
| *eat-2 spin-1* | 138 | 26.32 | 0.27 |
| *Figure 4A-B* | | | |
| *N2* | 158 | 19.63 | 0.39 |
| *spin-1,2* | 127 | 22.45 | 0.3 |
| *spin-1,2,3* | 149 | 17.94 | 0.16 |
| *eat-2* | 177 | 27.89 | 0.39 |
| *eat-2 spin-1,2* | 104 | 24.23 | 0.49 |
| *eat-2 spin-1,2,3* | 126 | 20.78 | 0.3 |
| *Figure 4E* | | | |
| *N2* | 161 | 19.98 | 0.43 |
| *SVIP-OE* | 170 | 19.26 | 0.37 |

C

| condition | χ2 | p-value | Corrected p-value |
|---|---|---|---|
| *Figure S6A* | | | |
| *eat-2 v.s. eat-2 spin-1* | 19.61 | 0.0000095 | 0.000038 |
| *Figure 4A-B* | | | |
| *N2* v.s. *eat-2* | 157.41 | 0 | 0 |
| *N2* v.s. *spin-1,2* | 12.84 | 0.0003 | 0.0017 |
| *N2* v.s. *spin-1,2,3* | 27.27 | 1.80E-07 | 8.90E-07 |
| *N2* v.s. *eat-2 spin-1,2* | 38.81 | 0 | 0 |
| *N2* v.s. *eat-2 spin-1,2,3* | 0.99 | 0.3204 | 1 |
| *eat-2* v.s. *spin-1,2,3* | 303.12 | 0 | 0 |
| *eat-2* v.s. *spin-1,2* | 107.62 | 0 | 0 |
| *eat-2* v.s. *eat-2 spin-1,2* | 29.15 | 6.70E-08 | 3.30E-07 |
| *eat-2* v.s. *eat-2 spin-1,2,3* | 157.77 | 0 | 0 |
| *spin-1,2* v.s. *spin-1,2* | 139.81 | 0 | 0 |
| *spin-1,2* v.s. *eat-2 spin-1,2* | 13.37 | 0.0003 | 0.0013 |
| *spin-1,2* v.s. *eat-2 spin-1,2,3* | 11.69 | 0.0006 | 0.0031 |
| *spin-1,2,3* v.s. *eat-2 spin-1,2* | 140.89 | 0 | 0 |
| *spin-1,2,3* v.s. *eat-2 spin-1,2,3* | 71 | 0 | 0 |
| *eat-2 spin-1,2* v.s. *eat-2 spin-1,2,3* | 34.23 | 4.90E-09 | 2.40E-08 |
| *Figure 4E* | | | |
| *N2* v.s. *SVIP-OE* | 2.93 | 0.0867 | 0.0867 |

**Fig. S6, related to Fig. 4: Statistical analyses of lifespans.**
(**A**) Lifespan of *WT*, *spin-1,2* and *spin-1,2,3* mutants. (**B**) Descriptive statistics for lifespans in Figure 4A-B, E and S6A. (**C**) Log-rank test results for lifespan comparisons in Figure 4A-B, E and S6A.

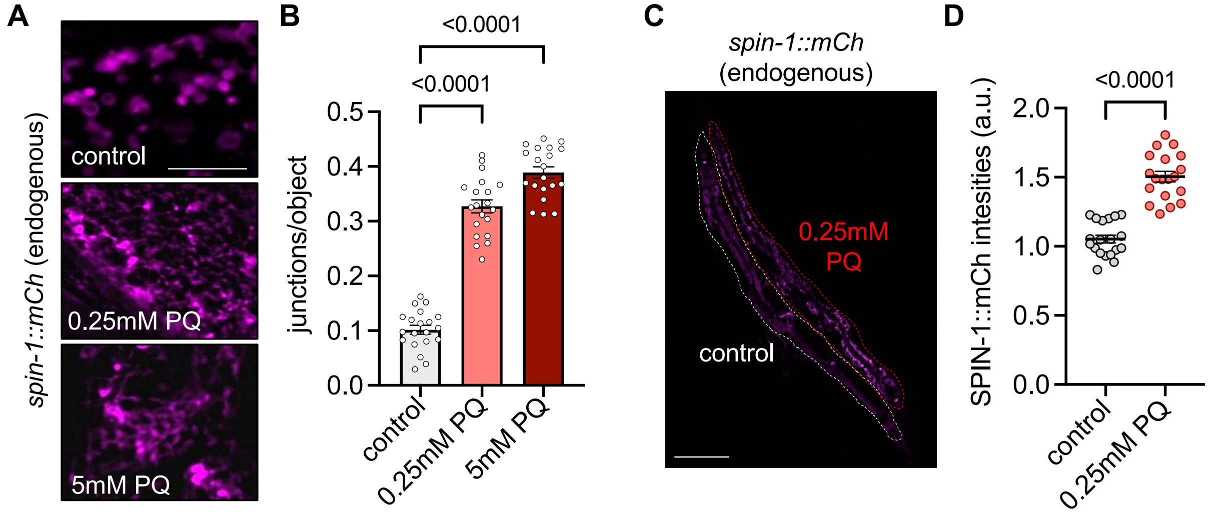


**Fig. S7, related to Fig. 4: Oxidative stress induces TLs in the *C. elegans* gut.**
(**A**) Representative images of endogenously-tagged *spin-1::mCh* in worms treated with control (0mM), low (0.25mM) or high (5mM) doses of Paraquat (PQ). Bar, 5µm. (**B**) Quantification of lysosomal junctions/object. Statistical significance was determined using one-way ANOVA with Dunnett's multiple comparisons (p-values indicated on graph). (**C**) Representative images of *spin-1::mCh* expression levels in worms treated with control or 0.25mM PQ. Bar, 100µm. (**D**) Quantification of SPIN-1::mCh fluorescence intensities. Statistical significance was determined using student's t-test (p-values indicated on graph).

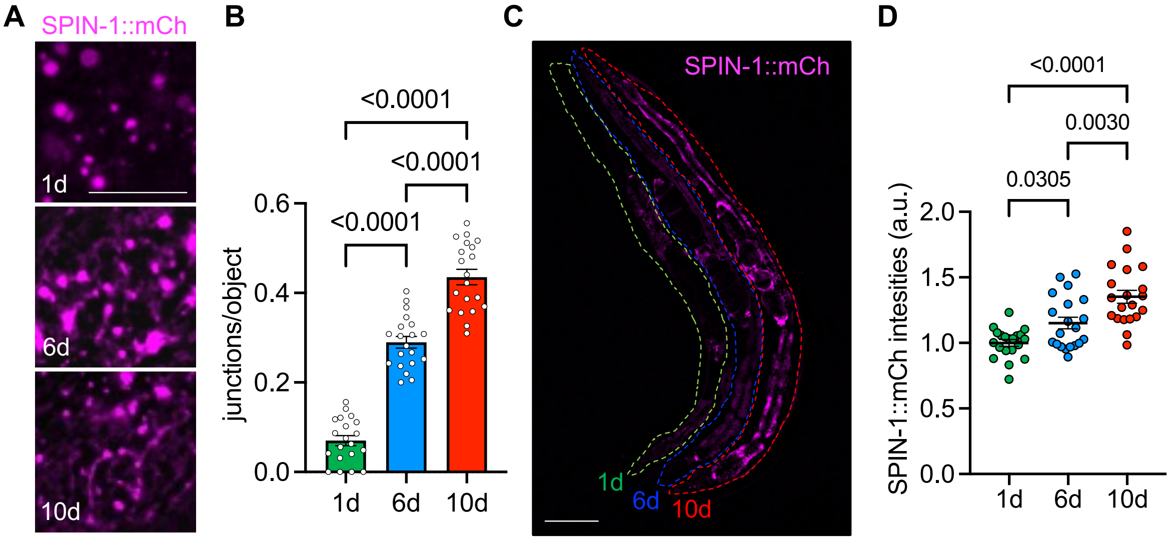


**Fig. S8, related to Fig. 4: Natural aging induces TLs in the *C. elegans* gut.**
(**A**) Representative images of endogenously-tagged *spin-1::mCh* in adult worms at 1, 6 and 10 days of adulthood. Bar, 5µm. (**B**) Quantification of lysosomal junctions/object. (**C**) Representative images of *spin-1::mCh* expression levels in adult worms at 1, 6 and 10 days of adulthood. Bar, 100µm. (**D**) Quantification of SPIN-1::mCh fluorescence intensities. Statistical significance was determined using one-way ANOVA with Dunnett's multiple comparisons (p-values indicated on graphs).

| Strain name | Genotype | Source |
|---|---|---|
| DA1116 | *eat-2(ad1116)* II | CGC |
| RB1678 | *spin-1(ok2087)* V | CGC |
| RB1702 | *spin-2(ok2121)* IV | CGC |
| RB1778 | *spin-3(ok2286)* X | CGC |
| COP2331 | *spin-1(knu1010[spin-1::mCherry::loxP::HygR::loxP])* V | this study (by In Vivo Biosystems) |
| COP2341 | *spin-2(knu1018[spin-2::mCherry::loxP::HygR::loxP])* IV | this study (by In Vivo Biosystems) |
| COP2343 | *spin-3(knu1020[spin-3::mCherry::loxP::HygR::loxP])* X | this study (by In Vivo Biosystems) |
| KAB3 | *louEx3(Pges-1::mCherry-gfp::lgg-1::unc-54 UTR)* | this study |
| KAB12 | *louEx12(Pges-1::svip::gfp::unc-54 UTR + Pges-1::spin-1::mCherry::unc-54 UTR)* | this study |
| KAB37 | *louIs4(Pges-1::spin-1::mCherry::unc-54 UTR)* X | this study |
| KAB58 | *louIs5(Pges-1::svip::unc-54 UTR + Podr-1::rfp)* | this study |
| KAB62 | *louIs5(Pges-1::svip::unc-54 UTR + Podr-1::rfp)*; *spin-1(knu1010[spin-1::mCherry::loxP::HygR::loxP])* V | this study |
| KAB63 | *eat-2(ad1116)* II; *spin-1(knu1010[spin-1::mCherry::loxP::HygR::loxP])* V | this study |
| KAB67 | *spin-2(ok2121)* IV; *spin-1(ok2087)* V | this study |
| KAB69 | *eat-2(ad1116)* II; *spin-1(ok2087)* V | this study |
| KAB70 | *eat-2(ad1116)* II; *spin-2(ok2121)* IV; *spin-1(ok2087)* V | this study |
| KAB72 | *spin-2(ok2121)* IV; *spin-1(ok2087)* V; *spin-3(ok2286)* X | this study |
| KAB90 | *eat-2(ad1116)* II; *spin-2(ok2121)* IV; *spin-1(ok2087)* V; *spin-3(ok2286)* X | this study |

**Table S1.** Strains used in this study.

**Movie S1.**

Representative movie demonstrating thrashing rates of WT and SVIP gut over-expression worms (SVIP-OE) at day 7 of adulthood.

**Movie S2.**

Representative movie demonstrating thrashing rates of WT and SVIP gut over-expression worms (SVIP-OE) at day 10 of adulthood.
